# *ZmCOL3*, a CCT-domain containing gene affects maize adaptation as a repressor and upstream of *ZmCCT*

**DOI:** 10.1101/205237

**Authors:** Minliang Jin, Xiangguo Liu, Wei Jia, Haijun Liu, Wenqiang Li, Yong Peng, Yanfang Du, Yuebin Wang, Yuejia Yin, Xuehai Zhang, Qing Liu, Min Deng, Nan Li, Xiyan Cui, Dongyun Hao, Jianbing Yan

## Abstract

Flowering time is a vital trait to control the adaptation of flowering plants to different environments. CCT-domain containing genes are considered to play an important role in plants flowering. Among 53 maize CCT family genes, 28 of them were located in the flowering time QTL regions and 16 genes were significant associated with flowering time based on candidate gene-based association mapping analysis. Furthermore, a CCT gene named as *ZmCOL3* was validated to be a flowering repressor upstream of *ZmCCT* which is one of the key genes regulating maize flowering. The overexpressed *ZmCOL3* could delay flowering time about 4 days whether in long day or short day conditions. The absent of one cytosine in 3’UTR and the present of 551bp fragment in promoter regions are likely the causal polymorphisms which may contribute to the maize adaptation from tropical to temperate regions. *ZmCOL3* could transactivate *ZmCCT* transcription or interfere circadian clock to inhibit flowering which was integrated in the modified model of maize photoperiod pathway.

**Highlight:** Maize CCT genes influence flowering time in different latitude environments and one of them named *ZmCOL3* is a flowering time repressor which could transactivate *ZmCCT* transcription to delay flowering.

## Introduction

The transition from vegetative to reproductive growth is a key developmental switch in flowering plants (Bluemel *et al.*, 2015). In seed crops, the switch could determine the production of dry matter (Jung and Müller, 2009). Understanding the molecular mechanisms underlying flowering is crucial for crop improvement. Maize is one of the most important crops globally for food, feed, and fuel. Domesticated from teosinte (*Zea mays ssp. parviglumis*) about 10,000 years ago in southwestern Mexico (Matsuoka *et al.*, 2002), maize was spread around the world from its tropical geographic origin with continuous improvements. It has been estimated that less than 1,000 genes were involved in the maize adaptation process (Liu *et al.*, 2015). Today, some tropical lines still flower late or fail to flower in temperate regions, limiting the utilization of tropical germplasm resources. Previous studies have shown that differences in maize flowering time were caused by the cumulative effects of numerous small effect quantitative trait loci (Buckler *et al.*, 2009). Recently, 90 flowering time regions were identified and 220 candidate genes were proposed through the use of a large population with nearly one million single-nucleotide polymorphisms (SNPs, Li *et al.*, 2016). A few key flowering genes, such as *Vgt1* (Salvi *et al.*, 2007) and *ZmCCT* (Huang *et* al., 2012; Yang *et al.*, 2013), have been cloned. However, little is known about the molecular mechanism and regulation pathway of flowering time in maize.

The basic genetic components of genes controlling flowering have largely been conserved in plants (Dong *et al.*, 2012). A large number of flowering genes in *Arabidopsis thaliana* and *Oryza sativa* have been well studied. The cloned genes participate in two main pathways: the circadian clock-controlled flowering and the photoperiod regulation of flowering. In *Arabidopsis*, the central oscillator of circadian clock consists of a series of negative feedback loops. *TOC1*, *CCA1,* and *LHY* are involved in the central loop. *CCA1* and *LHY* negatively regulate *TOC1* transcription in daylight and *TOC1* functions as a transcriptional repressor to *CCA1* and *LHY* at night (Alabadí *et al.*, 2001; Gendron *et al.*, 2012). The morning loop includes *CCA1, LHY,* and some PRR genes, such as *PRR7* and *PRR9* (Farré *et al.*, 2005; Nakamichi *et al.*, 2010). *GI* is the output signal of the circadian clock and it regulates *CO* transcription (Fowler *et al.*, 1999; Park *et al.*, 1999). *CO* up regulates *FT* expression, resulting in early flowering in long day conditions (Putterill *et al.*, 1995; Kardailsky *et al.*, 1999; Onouchi *et al.*, 2000). The majority of circadian clock genes are conserved in rice, including the *GI-CO-FT* pathway. *Hd1*, the *CO* homolog gene in rice, has a dual function. It promotes flowering in short day conditions but delays flowering in long day conditions by regulating *Hd3a*, the *FT* ortholog gene (Yano *et al.*, 2000; Kojima *et al.*, 2002; Hayama *et al.*, 2003). In addition, the *OsGI-Ehd1-Hd3a* flowering pathway is specific to rice as compared with *Arabidopsis. Ehd1* positively regulates *Hd3a* transcription and accelerates flowering in long day conditions (Doi *et al.*, 2004; Itoh *et al.*, 2010). Another rice-specific flowering gene, *Ghd7*, repress *Ehd1* expression to reduce the expression of *Hd3a* thus delaying the heading date of rice (Xue *et al.*, 2008).

*CO* contains the CCT (CO, CO-LIKE, and TIMING OF CAB1) domain, which is a conserved module with ~43 amino acids near the C terminus of proteins (Putterill *et al.*, 1995; Strayer *et al.*, 2000; Robson *et al.*, 2001). CCT domain-containing genes (CCT genes) play important roles in the circadian clock and the photoperiod flowering pathway. In addition to *CO*, *TOC1*, *PRR5*, and *PRR7* also are CCT domain genes. In total, 41 CCT genes were identified in rice and 11 of them have been determined to affect the flowering time, including *Hd1* and *Ghd7* (Yano *et al.*, 2000; Xue *et al.*, 2008; Zhang *et al.*, 2015). In maize, two CCT genes (*CONZ1* and *ZmCCT*) were also identified to affect flowering time. These studies underline the considerable importance of CCT genes in regulating plant flowering time.

The conservation of regulatory pathways and genes provides an excellent opportunity to systematically study maize flowering time regulation based on information from model species such as rice and *Arabidopsis*. In maize, CCT genes have not been identified and their association with flowering time has not been extensively explored. In the present research, we set out to identify the CCT genes in the maize genome and study their functions by comparing the identified genes’ locations in the genome with the flowering time related QTLs and evaluate the associations between natural variations of each CCT gene with flowering time by using candidate gene association studies. Moreover, the functions of a strong candidate gene, *ZmCOL3*, were validated with multiple approaches and it was added to the modified maize photoperiod pathway.

## Materials and methods

### Identification of CCT genes in maize

The HMM file of the CCT domain was downloaded from the PFAM dataset (http://pfam.xfam.org/, PF06203). HMMER software (Finn *et al.*, 2011), which uses probabilistic hidden Markov models (profile HMMs), was used to search sequences containing the CCT domain in maize genomes (B73_V2, Schnable *et al.*, 2009). The alignments of CCT protein sequences were done by Clustalx2.1 (Larkin *et al.*, 2007) and the unrooted polygenetic tree was constructed with MEGA 5.05 (Tamura *et al.*, 2011) by the neighbor-joining method and bootstrap analysis (1,000 replicates). The tree figure and main protein domain distributions were drawn by GSDS (http://gsds.cbi.pku.edu.cn/). The protein sequences of *Oryza sativa*, *Sorghum bicolor*, and *Brachypodium distachyon* were downloaded from RGAP (http://rice.plantbiology.msu.edu/index.shtml) and Ensemblplants (http://plants.ensembl.org/index.html). The same identification and polygenetic analysis methods were performed in these three species.

The expression pattern heatmap of maize CCT family members was drawn by the R language, and expression data came from PLEXdb (http://www.plexdb.org/).

### Candidate gene association mapping and homolog analysis

The polymorphisms of the identified CCT genes were extracted from a diverse association mapping panel contain 368 inbred lines and genotyped with more than 0.5 million SNPs (Li *et al.*, 2013; Fu *et al.*, 2013). Finally, 821 SNPs from 45 CCT genes including the 2Kb upstream/downstream regions were obtained. The association mapping panel was grown in 13 environments (**Table S1**). Flowering time was investigated and measured as days to tassel (DT), days to anthesis (DA), and days to silking (DS). A mixed linear model, which accounted for population structure and relative kinship (Zhang *et al.*, 2010), implemented in TASSEL3.0 software (Bradbury *et al.*, 2007) was used to identify the loci significant for flowering time.

The nucleotide sequences of *Oryza sativa* downloaded from RGAP (http://rice.plantbiology.msu.edu/index.shtml) and B73_RefGen_v2 were used to identify genome synteny alignments by SyMap V4.2 (Soderlund *et al.*, 2006). The homolog relationships between maize CCT genes and rice genes were integrated by using synteny blocks information and grass synteny orthologs provided by a previous study (Schnable *et al.*, 2012).

Maize QTL were obtained from Li’s study (Li *et al.*, 2016) and the homolog relationships between rice known genes and maize CCT genes were drawn by Circos-0.62-1 (http://circos.ca/).

### Subcellular localization of *ZmCOL3*

The full-length *ZmCOL3* coding sequence (T01; 1,008bp) was amplified from B73 cDNA and cloned downstream of the CaMV35S promoter in the pCAMBIA1302 vector that carried GFP. The nuclear localized gene *HY5* in Arabidopsis (Chattopadhyay *et al.*, 1998) was inserted into pCAMBIA1301 vector that carried RFP and was used as a nuclear marker. Both constructs were introduced into maize protoplasts as described (Yoo *et al.*, 2007). The GFP and RFP signals were detected using FV1200 Laser Scanning Microscope (OLYMPUS CORPORATION).

### Transformation of *ZmCOL3*

The 1,114bp DNA fragment of *ZmCOL3* from ATG to TGA (refer to B73) was synthetized and used for the transformation study. The ZmUbi promoter driving this sequence was inserted into the modified binary vector pCAMBIA3300, which contained the selectable marker PAT coding region driven by 2× CaMV35S promoter, TEV intron, and vsp terminator sequences. Immature zygotic embryos of maize hybrid, HiII (B73 × A188) were infected with *A. tumefaciens* strain EHA105 harboring the binary vector based on the published method (Frame *et al.*, 2002). In brief, F_2_ immature zygotic embryos (1.5–2.0 mm) of the maize HiII hybrid genotype were excised from a maize ear harvested 10 to 13 days after pollination. After co-culturing the excised immature embryos with Agrobacterium carrying the targeted vector, the immature embryos were placed on selection medium containing bialaphos (3 mg /L) and cefotaxime (250 mg /L) in order to inhibit the growth of untransformed plant cells and excess Agrobacterium. Putatively transformed events were identified as early as 5 weeks after infection. Regeneration of R0 transgenic plants from Type II embryogenic callus was accomplished by a 2- to 3-week maturation step on Regeneration Medium I, followed by germination in the light on Regeneration Medium II. Regenerated R0 plantlets were moved to soil, where they were sampled and grown to maturity in greenhouse conditions. The following detailed breeding information can be seen in **Fig. S1**. Transgenic plants were identified by herbicide test, bar test papers, and PCR validation of bar and *ZmCOL3* (primer BAR and COL3 in **Table S2**). Leaf tissues were collected from plants at V6 stage (Vegetative 6, six fully extended leaves). RNA of each sample was isolated by a quick RNA isolation kit (HUAYUEYANG, Beijing, China). cDNA was synthesized using one-step gDNA removal and cDNA synthesis supermix (TransGen, Beijing, China). *ZmCOL3* expression was identified via real-time PCR by primer qCOL3 and qCOL3-2 (TransStart® Tip Green qPCR SuperMix from TransGen, Beijing, China), using primer ACTIN as the internal control. The receptor HiII was used as a control to evaluate relative gene expression levels of transgenic plants.

In total, 30 ears were randomly chosen from each transgenic event including 15 positive transgenic lines and 15 negative controls. Same type ears were measured for weight related traits as a whole (cob weight, kernel weight), and values were divided by ear number. Other traits (ear length, ear diameter, ear row number, cob diameter, and kernel number per row) were tested individually. After threshing, 50 kernels from each ear type were randomly picked out to measure kernel shapes (length, width and thickness). The measurements of about 100 kernel weight and volume were repeated three times and the average value was obtained.

### Re-sequencing of *ZmCOL3*

The primers (**Table S2**) were used to amplify and sequence the promoter (~1.6kb) and gene region of *ZmCOL3* in 152 maize lines. The sequences were aligned by ClustalX2.1 (Larkin *et al.,* 2007), SNP-sites (https://github.com/sanger-pathogens/snp-sites) was used for the calling of SNPs and InDels. Association mapping analysis was performed with Tassel3.0 software (Bradbury *et al.*, 2007) and the levels of LD between pairs of sites were calculated using Haploview (Barrett *et al.*, 2005).

### RNA sequencing of transgenic event 1-39

T1 plants of transgenic event 1-39 in 2016JL and T3 plants of 1-39 in 2016DHN were used to perform RNA sequencing. Leaf tissues were harvested in V3 (Vegetative 3, three fully extended leaves) and V6 stages for T1 plants and V6 stages for T3 plants. RNAs of 10 transgenic plants and 10 controls were mixed as one sample, respectively. Three technology replicates were performed for each sample. In sum, 18 samples were used for sequencing. RNA libraries were constructed according to TruSeq® Stranded mRNA Sample Preparation Guide (TruSeq® Stranded mRNA LT-SetA. RS-122-1201). The size of each prepared library ranged between 200bp to 500bp long. All libraries were PE150 sequenced using HiSeq3000. Low quality reads were filtered by seqtk_1.0 (https://github.com/lh3/seqtk) and trimmomatic-0.33 (Bolger *et al.*, 20140). STAR-2.5.2b (Dobin *et al.*, 2013) was used to align RNA-seq reads to the reference genome. Differentially expression was analyzed by an R package DESeq2 (Love *et al.*, 2014). The primers (**Table S2)** were used to measure expression in individual by qRT-PCR (10 controls and 10 transgenic lines in V6 stage of T1 Plants).

### Transient assays for *in vivo* activation activity

The reporters were constructed based on pGreenII 0800-LUC vector (Hellens *et al.*, 2005) and the effectors were constructed based on the pRI 101-AN vector. The promoter fragments of *ZmCCT* (−643 to −1 bp) and *ZCN8* (−2,000 to −1 bp) were amplified from B73 genomic DNA by PCR (**Table S2)**. To clone into pGreenII 0800-LUC vector, the HindIII site was used. And the full length of *ZmCOL3* CDS (1,008bp) was amplified from B73 cDNA (**Table S2)**. It was cloned into the pRI 101-AN vector using SmaI site. Transient dual-luciferase assays in maize protoplasts were performed and measured using dual-luciferase assay reagents (PROMEGA CORPORATION) by PerkinElmer EnSpire. Five independent measures were carried out for each analysis.

## Results

### Identification of maize CCT genes

In total, 53 CCT genes were identified in the maize genome (**Table S3**), which were not randomly distributed on maize 10 chromosomes (**Fig. S2**). There were 11 CCT genes located on chromosome 5, while only 3 genes were situated on chromosomes 3, 7 and 8 respectively. An unrooted polygenetic tree was constructed based on the full length protein sequences of maize CCT genes (Fig. 1A). CCT genes can be divided into 4 clans: COL-like genes with CCT and one or two B-box type zinc finger domains, CMF-like genes with only CCT domain, PRR-like genes with CCT and response regulator receiver domains, TIFY genes with CCT, TIFY, and GATA zinc finger domains. The 11 genes that only contained CCT domain were clustered by sequence similarity into COL-like and PRR-like genes clans (8 in COL and 3 in PRR), and may have lost of B-box type zinc finger domain, the conserved response regulator receiver domain. We still considered them as COL- or PRR-like genes in this study. Finally, there were 27 COL-like, 15 CMF-like, 8 PRR-like, and 3 TIFY-like genes in maize. To further study the evolutionary relationship of CCT genes, 113 CCT gene sequences from other Poaceae plants (41 in *Oryza sativa*, 35 in *Sorghum bicolor* and 37 in *Brachypodium distachyon*, **Table S4**) were collected. Only the protein sequences of their highly conserved CCT domain were used for polygenetic analysis (**Fig. S3**), and were clearly separated into four subfamilies on this basis. COL-like genes were divided into three clans, which implies that their origin was different and that they may have different functional features. This is consistent with the suggestion that CCT genes were present and differentiated prior to the appearance of dicots since CCT genes were not clustered based on species (Cockram *et al.*, 2012).

**Fig. 1.**
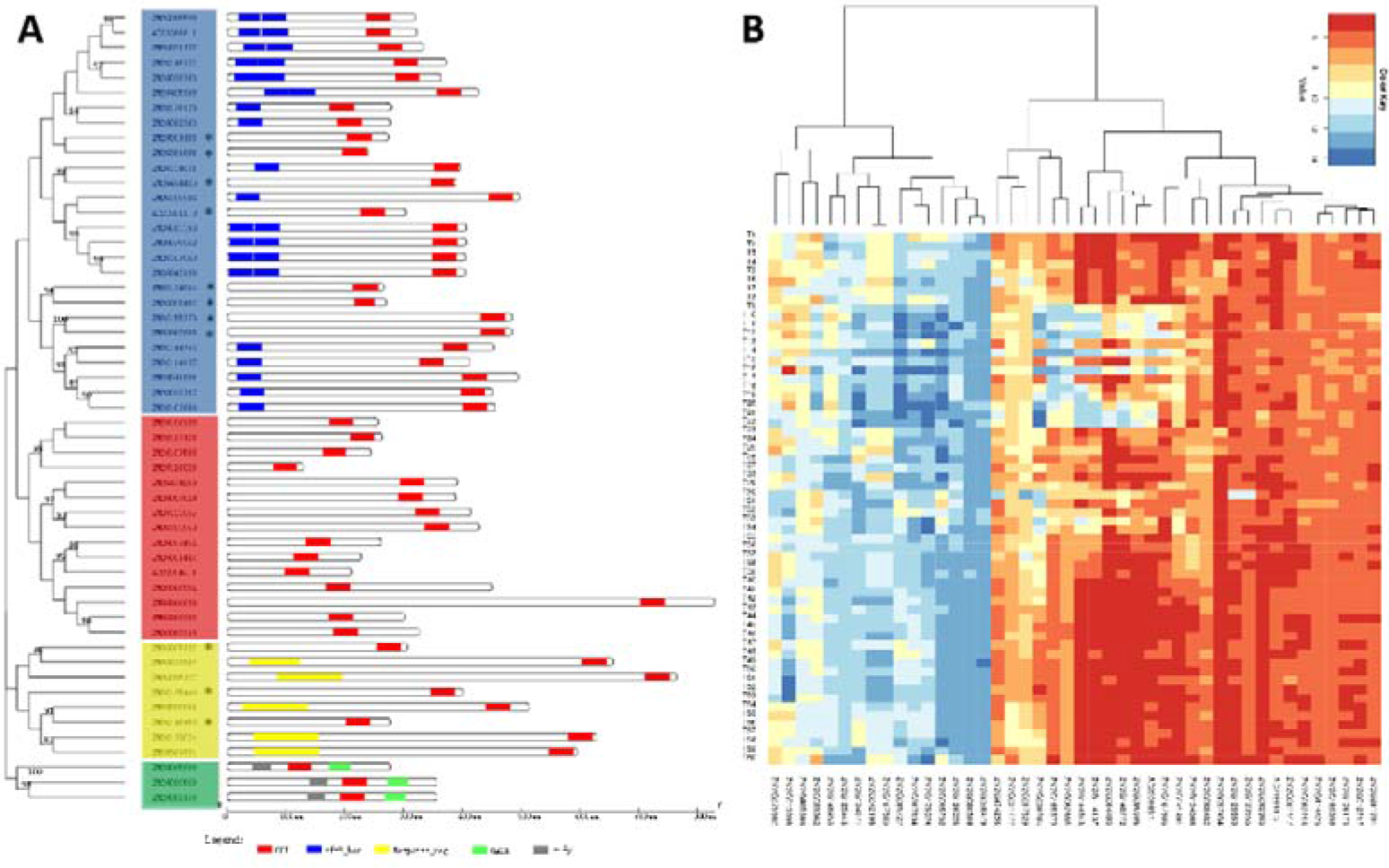
Polygenetic tree, gene structure, and expression pattern of maize CCT genes. (**A**) The neighbor-joining polygenetic tree of maize CCT genes was constructed based on full-length protein sequences. Bootstrap values from 1,000 replicates were indicated at each node (The values below 80 were not shown). The main protein domain distributions are shown after gene names. Maize CCT genes can be divided into four clans: COL-like genes (blue panel), CMF-like genes (red panel), PRR-like genes (yellow panel), and TIFY-like genes (green panel). (**B**) The expression pattern of 44 maize CCT genes based on B73 expression data of different tissues (Sekhon *et al.*, 2011). The degree from high to low expression level is shown from blue to red. T1-T3: Root; T4-T7: SAM and young stem; T8-T9: Whole seeding; T10-T22: Leaves; T23-T25: Internodes; T26-T27: Cob; T28-T30: Tassel and Anthers; T31: Silk; T32-T34: Husk; T35-T59: Seeds; T60: Germinating seed.

### Expression pattern of maize CCT genes

A comprehensive atlas of transcription pattern in 60 distinct tissues representing
11 major organ systems of maize inbred line B73 provided us an opportunity to observe the transcription characteristics of maize CCT genes (Sekhon *et al.*, 2011). The expression data for 44 CCT genes were obtained and could be divided into roughly two types: high expression genes (16) and low expression genes (28) (Fig. 1B). All of the PRR-like and TIFY-like genes expressed highly, while majority of CMF-like genes, except one, expressed lowly. For COL-like genes, there were 6 expressed highly and 21 expressed at low level.

### Maize CCT genes are associated with flowering time

To explore the associations between maize CCT genes and flowering time, we first compared the locations of identified CCT genes and mapped flowering time QTLs. In recent large-scale maize flowering time QTL mapping experiments, 130 flowering time-related QTLs were identified (Li *et al.*, 2016). Among 53 maize CCT genes, 28 were located in these QTL regions (28/53, 52.8%; Fig. 2A, **Table S5**), including 14 COL-like, 5 CMF-like, 7 PRR-like, and 2 TIFY-like genes.

**Fig. 2.**
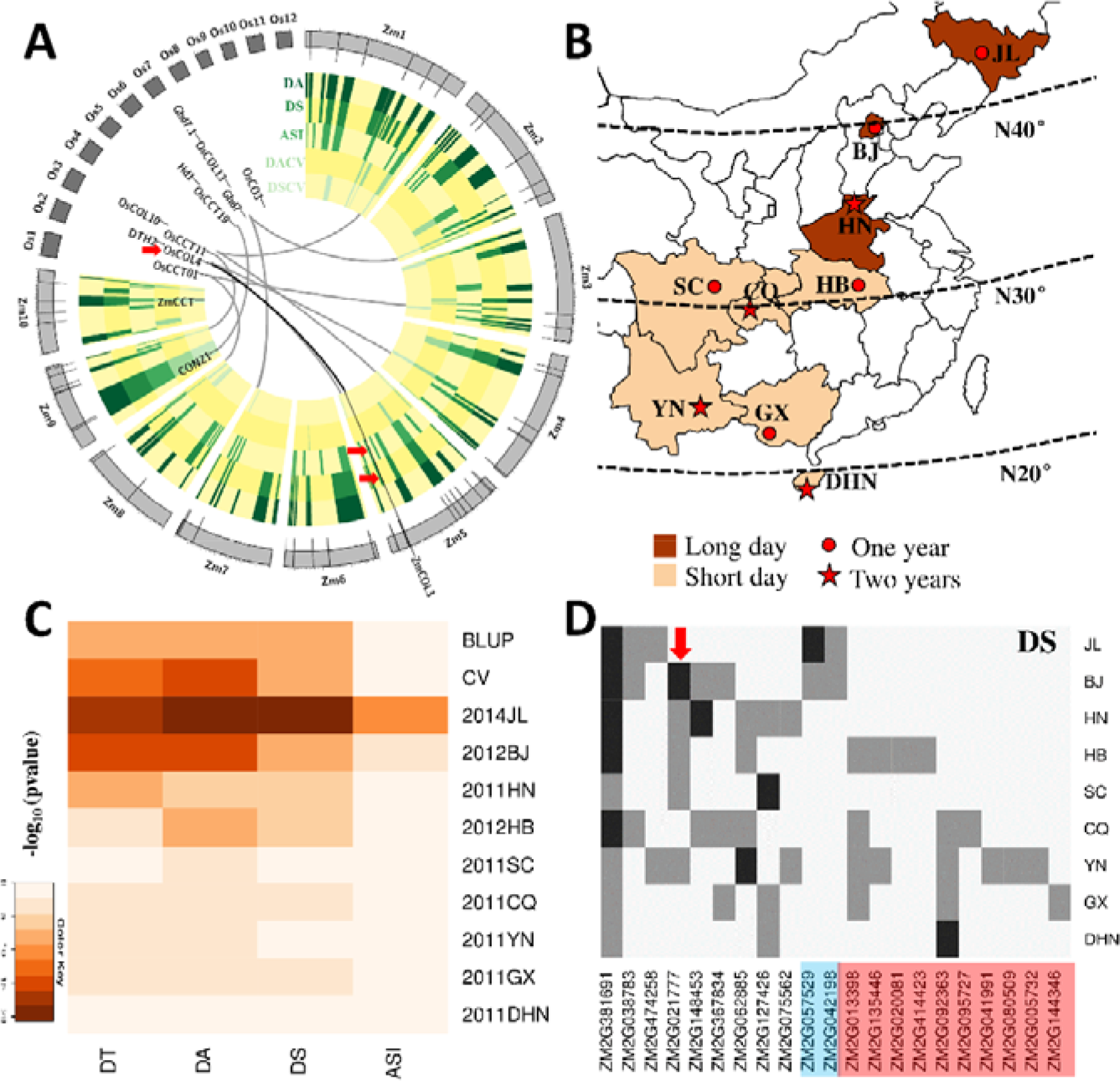
The associations of maize CCT genes and flowering time. (**A**) Comparative analysis of maize and rice CCT genes. The black lines on maize chromosomes represent the physical positions of maize CCT genes. Maize flowering time QTL regions are plotted around the maize genomes in green. From outside to inside the circles represent QTLs for the following traits: DA (Days to Tassel), DS (Days to silking), ASI (Anthesis-Silking Interval), DACV and DSCV (The coefficients of variations for DA and DS). The links in the circle are the homolog relationship between maize CCT genes and rice known heading date genes. (**B**) Mapping population panel phenotyping site locations in China (detailed information in **Table S1**). (**C**) The size of the phenotypic effect of *ZmCCT* in different environments based on the candidate gene association analysis. (**D**) Candidate gene association mapping results of DS in different locations. The phenotypic effect of HN (Henan; E114°, N36°), CQ (Chongqing; E107°, N30°), YN (Yunnan; E103°, N23°), and DHN (Hainan; E109°, N18°) in two years were merged using the average values. Grey blocks represent significance at P<0.01 and black blocks represent significance at P<0.001. The genes shown in blue were only detected in long day conditions and the genes shown in red were found in short day conditions.

Candidate genes association mapping analysis was performed for all of the maize CCT genes. In total, 821 SNPs of 45 CCT genes were obtained from the high-density genotype data in 368 diverse inbred lines called AMP (Association Mapping Population Panel, Fu *et al.*, 2013), which were phenotyped in 13 environments (Fig. 2B, **Table S1**) for DT (Days to Tassel), DA (Days to Anthesis) and DS (Days to Silking). The difference between DA and DS was estimated as the anthesis-silking interval (ASI). Average daylight time greater than 14 hrs in the two months after seeding was considered as long day conditions (4 environments), while less than 14 hrs was considered short day conditions (9 environments). The flowering times measured in multiple environments were analyzed in three ways. (1) The best linear unbiased predictions (BLUPs) were used to generalize the overall performance and used for association studies. The broad sense heritabilities of DT, DA, DS, and ASI of AMP across 13 environments reached 94.3%, 94.2%, 92.6%, and 67.1%, respectively. (2) Variation of flowering time in 13 environments was used to reflect the environmental response of the association mapping panel. The coefficients of variations (CV) for DT, DA, and DS of each line were estimated and used for association studies. These coefficients varied from 9.7% to 26.9% for DTCV, from 10.4% to 26.2% for DACV, and from 10.2% to 28.0% for DSCV. (3) Phenotype in each environment was used for association study to discover the potential environment-specific flowering effectors. In total, 34 and 16 CCT genes were found to significantly associate with flowering time at P<0.01 and 0.001, respectively (**Table S6**). *ZmCCT* was detected in multiple environments and the effect size increased with the increase in latitude, which is consistent with the previous study (Yang *et al.*, 2013, Fig. 2C). Some CCT genes were environmentally sensitive (**Fig. S4**; Fig. 2D), and only affect the flowering time in either long or short day conditions. For instance, GRMZM2G057529 was significantly associated with DT, DA, and DS only in long day conditions in JL (Jilin; E125°, N44°), while GRMZM2G092363 influences DT and DA in YN (Yunnan; E103°, N23°) and DHN (Hainan; E109°, N18°), which represent short day conditions. In addition, rice homologs of 13 maize CCT genes also influence heading data (Fig. 2A, **Table S7**). All of the above results provide credible evidence for the importance of CCT-like genes in the control of flowering time in maize.

### *ZmCOL3* is an important candidate for maize flowering time

Combining bioinformatics analysis, QTL mapping, and candidate gene association study (**Fig. S5**), a strong candidate, *ZmCOL3*, the homolog of rice flowering repressor *OsCOL4* (Lee *et al.*, 2010) was chosen for functional verification. *ZmCOL3* is located on chromosome 5 close to the QTL affecting DS and ASI. One SNP (SNP-1296) within the gene significantly associated with DS in BJ (Beijing; E116°, N40°; P=8.0E-04), and one SNP (SNP-1351) significantly associated with ASI in HN (Henan; E114°, N36°; P=7.3E-04). The difference of DS between two alleles in BJ is about 11 days. *ZmCOL3* appears to affect flowering time primarily in long day conditions as BJ and HN locations are in high latitude regions.

According to the atlas of transcription during different maize development stages (Sekhon *et al.*, 2011), *ZmCOL3* expressed higher in leaves, internodes, and embryo (**Fig. S6**). The time-course expression pattern of *ZmCOL3* in leaves indicated that it reached highest levels at 3.5 hrs after sunrise and then fell quickly (Fig. 3A). The vector that contained the CaMV35S promoter driving transcription of *ZmCOL3-GFP* was introduced into maize protoplasts. The GFP signal was detected in the nucleus (Fig. 3B), which is consistent with a previous finding before that the CCT domain could be functional in nuclear localization of proteins (Robson *et al.*, 2001). We generated transgenic plants overexpressing *ZmCOL3*, to further assess its function (**Fig. S7**). Four independent transgenic events were obtained. The transgenic lines and normal materials contained identical transcripts, which confirmed the accuracy of transformation (**Fig. S8**). The expression level of *ZmCOL3* in transgenic plants was over 10 times higher than the controls (Fig. 3C, **Fig. S9**). The families from T1 to T3 were planted in JL, at long day conditions, and DHN, at short day conditions. The transgenic plants exhibited significant phenotypic differences compared with controls in the long day condition (JL), with later flowering (by 4.0d, P=2.3E−121), higher plant and ear height (by 21.2cm and 17.1cm, P=1.2E−26 and 2.4E−39, respectively), and higher internode number (by 1.5, P=9.3E−64) (Fig. 3D, E, **Table S8**). However, there was not a simple positive correlation between flowering time and yield traits. For instance, volume weight and kernel length increased in transgenic lines when the flowering time difference between transgenic lines and controls reached 3d (1-39-T2) but decreased when transgenic lines flowered late by 5d (29-5-T1). Similar phenotypes also appeared in the short day condition (DHN), with later flowering by 3.2d (P=3.5E−63), higher plant and ear height by 10.0cm (P=4.4E−7) and 12.8cm (P=1.4E−33), and higher internode number by 1.6 (P=1.1E−46) (**Table S8**). It is worth noting that the background expression of *ZmCOL3* in DHN was extremely low. This suggests that *ZmCOL3* did not play a role in short day condition resulting in lack of significant associations with flowering time in short day condition. However, overexpressed *ZmCOL3* could influence flowering time in short day conditions. These results suggest that *ZmCOL3* may be photosensitive and function as a flowering repressor in maize.

**Fig. 3.**
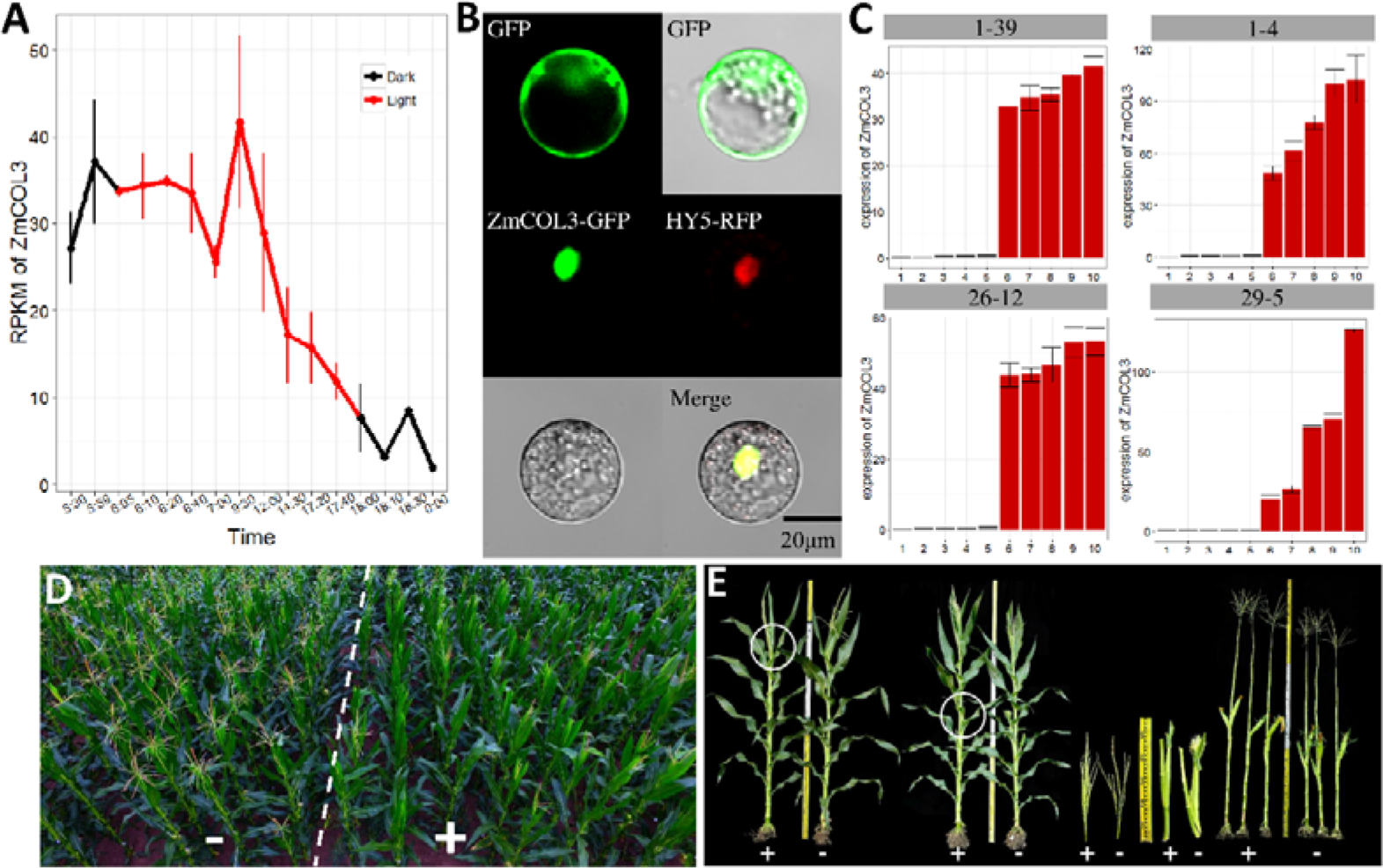
Functional validation of *ZmCOL3* by transformation analysis. (**A**) Diurnal rhythms of *ZmCOL3* expression in B73 leaves. Light was turned on at 6:00 and off at 18:00. (**B**) A pCAMBIA1302-35S:ZmCOL3-GFP construct was used to assess protein localization. The pCAMBIA1302-35S: GFP was used as a control and pCAMBIA1301-35S: HY5-RFP was used as a nuclear marker. ZmCOL3-GFP was detected in the nuclear of maize protoplast. (**C**) Expression of *ZmCOL3* in transgenic plants (red) and controls (grey) in 2016JL (Jilin; E125°, N44°). Error bars represent standard error (n = 3). (**D**) Field phenotype of transgenic event 1-4 in 2016JL. Transgenic plants (right, +) showed later tasseling than controls (left, −). The photo was taken at 68 days after planting. (**E**) The performance comparison in T1 generation of 1-39 between transgenic lines (+) and controls (−) in 2016JL. Transgenic lines flowered later (photo was taken at 70 days after planting), and had higher plant and ear height, and higher total internodes number (photo was taken at 87 days after planting).

We attempted to silence *ZmCOL3* transgenically. Its expression was inhibited, but not as much as expected (**Fig. S10**). The hairpin sequence introduced for silencing contained a short intron sequence to ensure specificity, which may have reduced silencing efficiency. The phenotype was consistent with the expectation that silenced transgenic lines would flower slightly earlier than controls (**Table S8**).

### Identification of the possible functional polymorphisms of *ZmCOL3*

In order to discover the underlying functional polymorphisms of *ZmCOL3*, 152 maize inbred lines from AMP were used for re-sequencing about 3Kb regions covering the promoter, coding, and 5’- and 3’- UTR regions. In total, 135 SNPs and 26 insertions and deletions (InDels) were discovered, including 7 non-synonymous SNPs and 3 InDels within exon regions resulting in the absence of amino acids rather than frameshift and premature termination. The calculation of pairwise linkage disequilibrium (LD) of *ZmCOL3* polymorphisms showed two clear and independent LD blocks (Fig. 4A). Two significant associations (PSNP-660 and InDel-3296) were detected affecting DS in BJ located in the two different LD blocks, which implied more than one functional polymorphism existed within *ZmCOL3* (Fig. 4A).

**Fig. 4.**
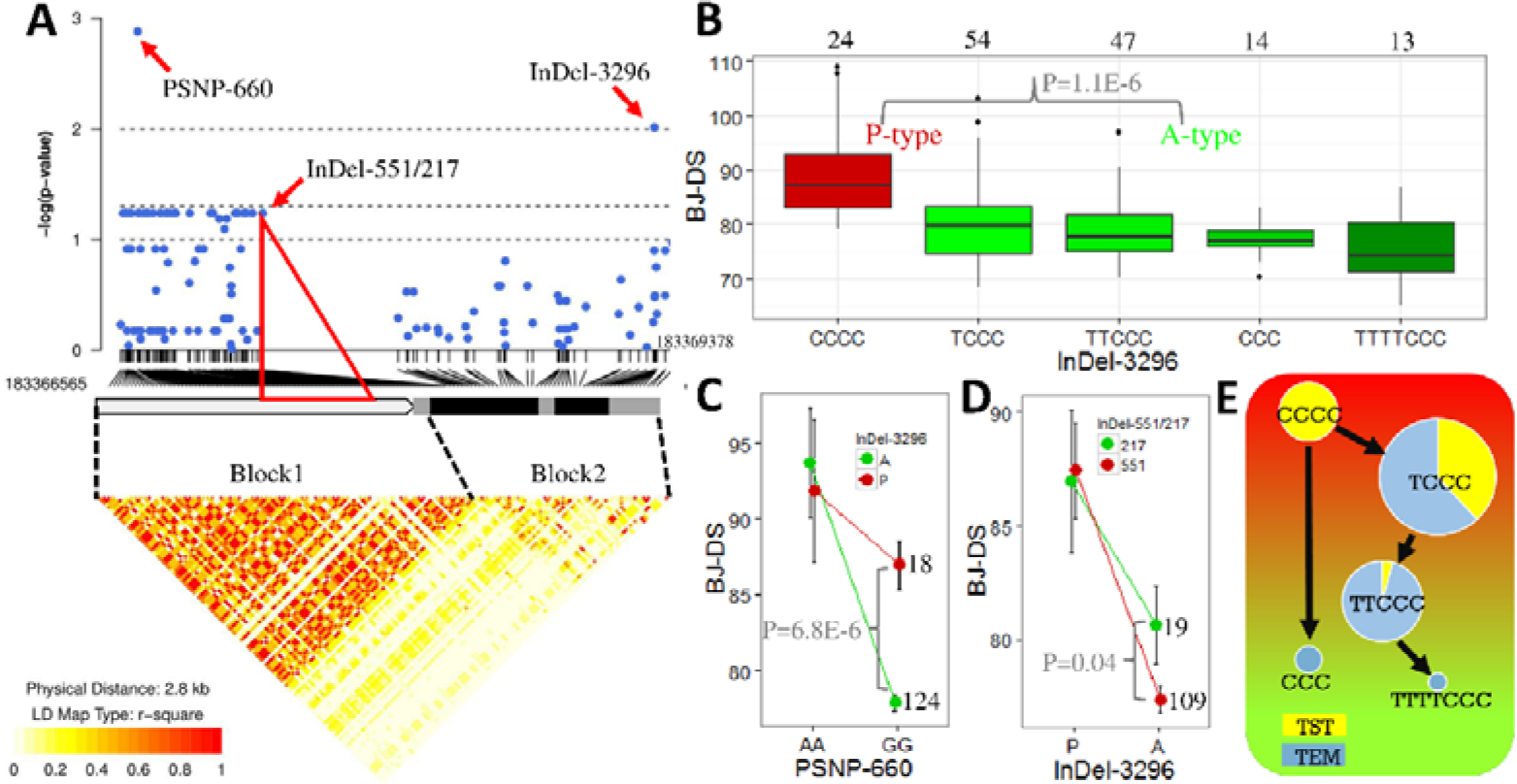
Possible functional sites in *ZmCOL3* gene. (**A**) Associations between polymorphisms within *ZmCOL3* and DS (Days to Silking) in BJ (Beijing; E116°, N40°) presented in plot figure. The gene structure is shown under the x-axis. White rectangle represents promoter region, grey rectangles represent UTR and intron, and black rectangles represent CDS region. The position of InDel-551/217 is indicated. The inverted triangle below gene structure indicate the locations of LD for polymorphisms within *ZmCOL3*. The polymorphisms with MAF<0.05 were excluded. (**B**) Allele analysis around InDel-3296. X-axis represents five allele combinations and y-axis shows the DS in BJ. Red boxes represent P-type and green boxplots are A-types. (**C**) Haplotype comparison of PSNP-660 and InDel-3296. The plot represents the median value of phenotypes. Error bars represent standard error. (**D**) Haplotype comparison of InDel-3296 and InDel-551/217. (**E**) Population distribution of InDel-3296. TST: tropical lines; TEM: temperate lines. Each circle represents an allele, and the size of the circle is proportional to the number of lines within the allele.

PSNP-660 is a rare allele located in Block1 that has few LD with other variations. InDel-3296 is located in the same position as SNP-1296, and is the most significant SNP by candidate gene association analysis. The TC (Thymine and Cytosine) enrichment around the loci makes it very difficult to distinguish each polymorphism within this region accurately and all five polymorphisms within this locus were combined together for haplotype analysis (Fig. 4B, **Table S9**). Five major haplotypes were identified. The haplotype containing four cytosine (P-type, CCCC) is associated with delayed flowering time compared with the other four haplotypes (A-types, Fig. 4B). We found that the effects of InDel-3296 and PSNP-660 were not independent, in that InDel-3296 affected flowering time more significantly (P=6.8E-6) when the allele of PSNP-660 was fixed as GG (Fig. 4C). However, the number of lines with the AA genotype was small (n=10) affecting the statistical calculation. In the promoter region, a 551bp insertion was present in some lines, while a totally different 217bp insertion exists in other lines at the same position. The InDel-551/217 is the common variant and the 551bp insertion was identified in 210 inbred lines among the 317 genotyped inbred lines, whereas the other 107 lines contain the 217bp insertion (**Table S10**). The InDel-551/217 variation was found to affect the flowering time significantly at JL, BJ, and HB (Hubei; E114°, N31°) locations (Student’s t test, P<0.01; **Table S11**). Interestingly, InDel-551/217 had different phenotypic performance when the allele of InDel-3296 was different, as indicate above (Fig. 4D).

In order to validate the genetic effects of InDel-3296 and InDel-551/217, tropical line CML189, which is P-type for InDel-3296 and 217bp insertion for InDel-551/217 (P-217-type), was crossed with the temperate line Mo17, which is A-type for InDel-3296 and 551bp insertion for InDel-551/217 (A-551-type), to generate a F_2_ population. The allele of PSNP-660 was monomorphic between in Mo17 and CML189. The F2 population was grown in three environments, including one in long day conditions (JL) and two in short day conditions (HB and DHN). About 300 individual plants were genotyped. As expected, a significant flowering time difference between homozygous P-217-type and homozygous A-551-type was observed in long day conditions (Student’s t test; DT, P=9.5E−3; DA, P=0.02; DS, P=8.0E−06, **Table S12**). No significant difference was detected in short day conditions, confirming that *ZmCOL3* affects flowering time primarily in long day conditions.

Among the 152 lines used for re-sequencing, there were 38 tropical lines and 66 temperate lines. The haplotype of the majority of the tropical lines was P-type (21 of 38, 55.3%; Fig. 4E), which was absent in temperate lines. All the temperate lines were A-types. Two of the A-types (CCC and TTTTCCC) were absent in tropical lines. We infer that three haplotypes were originally present in tropical lines (CCCC, TCCC and TTCCC) and that temperate lines retain two of them (TCCC and TTCCC) and the other two types (CCC and TTTTCCC) arose and were selected as adaptation to the long day conditions (Fig. 4E).

### A modified model of maize photoperiod pathway

In order to identify the downstream regulatory genes of *ZmCOL3*, RNA-seq data generated from transgenic plant 1-39 (see Materials and Methods) were obtained and used for differentially expressed genes (DEGs) analysis. Leaves from two stages, V3 and V6 in JL, were sequenced at the transcriptome level. High consistency was observed between three replications and the transcription differences between V3 and V6 stage were larger than between transgenic and control plants (Fig. 5A). As a transcriptional factor (TF), *ZmCOL3* regulates the differential expression of thousands of genes, including 2,072 genes expressed differently in both transgenic and control plants in two stages (padj≤0.01; Fig. 5B). 1,305 of them were present in GO database with annotation enriched in photosynthesis related terms (P<0.01, FDR<0.05; Fig. 5C). GRMZM2G005732 (*ZmPRR37a*) and GRMZM2G033962 (*ZmPRR37b*), the homologs of rice heading data repressor *OsPRR37* (Koo *et al.*, 2013), were regulated positively in transgenic lines. This was validated by qRT-PCR (Fig. 5D). In addition, *ZmCCT* was also confirmed to be up-regulated by *ZmCOL3* (Fig. 5D). *ZmCCT* was not detected by DEGs analysis, possibly because of its low level of expression. The expression of the florigenic gene *ZCN8* in controls was 1.5 fold higher than in transgenic lines, which is consistent with earlier flowering in controls, although the difference did not reach significance. *ZmCOL3* as a TF may bind to the promoter of downstream gene directly and affect its transcription. Luciferase (LUC)/Renillareniformis (REN) transactivation assays were carried out in maize protoplasts (Hellens *et al,* 2005) to test whether *ZmCOL3* can bind to the promoter of *ZmCCT* directly and increase its expression. Strong transactivation of LUC reporter gene expression was detected when it was driven by the 643bp promoter of *ZmCCT,* but not by the 2,000bp promoter of *ZCN8* (Fig. 5E). These results imply that *ZmCOL3* may directly increase *ZmCCT* expression, but not *ZCN8,* to influence flowering time in long day conditions.

**Fig. 5.**
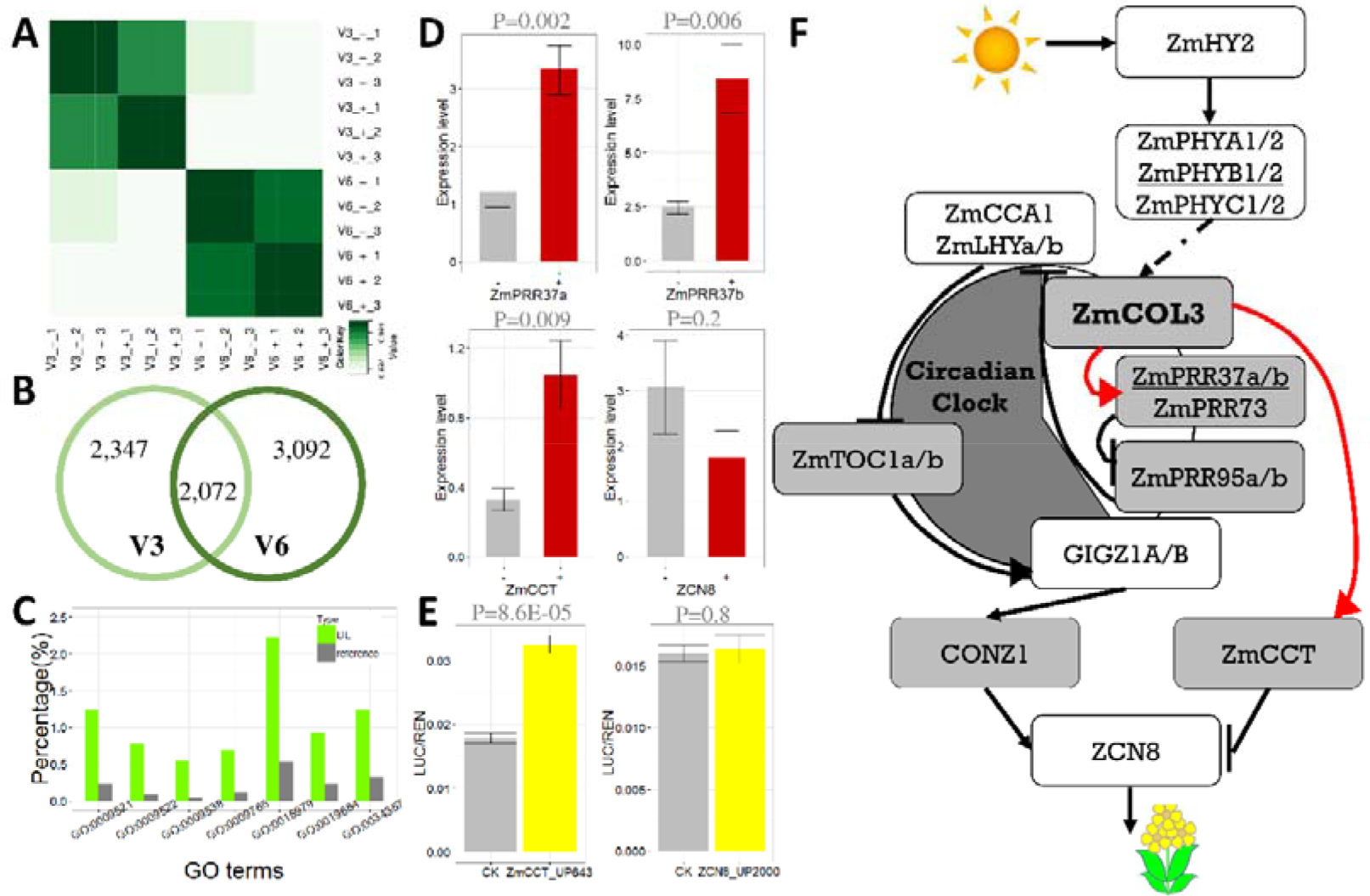
DEGs analysis of transgenic event 1-39 and a modified model of maize photoperiod pathway. (**A**) Expression correlation between 12 samples of transgenic event 1-39 in JL (Jilin; E125°, N44°). (**B**) Number of Differentially Expressed Genes (DEGs) at V3 and V6 in JL. (**C**) Enrichment in photosynthesis-related GO terms. The y-axis represents the percentage of genes belonging to each GO term. GO:0009521 photosystem; GO:0009522: photosystem I; GO:0009538: photosystem I reaction center; GO:0009765: photosynthesis, light harvesting; GO:0015979: photosynthesis; GO:0019684: photosynthesis, light reaction; GO:0034357: photosynthetic membrane. (**D**) RNA (+: transgenic lines; −: controls) of V6 in JL were used to evaluate expression of *ZmPRR37a/b*, *ZmCCT*, and *ZCN8*. Expression levels were normalized to actin. Error bars represent standard error (n=10). P-value of Student’s test is indicated. (**E**) Relative transactivation of *ZmCOL3* to *ZmCCT* and *ZCN8* in maize protoplast. LUC/REN indicates the ratio of the firefly luciferase activity and the Renilla luciferase activity. Error bars represent standard error (n = 5). CK represents samples with the reporter genes (LUC/REN) alone. (**F**) Proposed model for the *ZmCOL3* role in flowering pathway. Arrows indicate positive regulation and T-bars indicate suppression. The genes in grey blocks are maize CCT genes. The red lines were validated by qPCR or LUC/REN transactivation assays.

Since *ZmCOL3* mostly affects flowering time in long day conditions, the data from DNH (N18°) may be more suited to the DEG analysis than the data in JL (N44°). With the same standard (padj≤0.01), as many as 11,546 genes expressed differentially between transgenic and control lines. Besides *ZmPRR37a* and *ZmPRR37b*, some other genes participating in the circadian clock pathway were also directly or indirectly regulated by *ZmCOL3*, such as *ZmCCA1* (up-regulated), *ZmTOC1* (down-regulated) (**Table S13**). Combining these results with previous studies (Dong *et al.*, 2012, Yang *et al.*, 2013), we propose a modified model of the maize photoperiod pathway (Fig. 5F). We propose that *ZmCOL3* itself or genes upstream of *ZmCOL3* can sense the day length such that *ZmCOL3* is inhibited under short day conditions and activated in long day conditions. On the one hand, *ZmCOL3* could transactivate *ZmCCT* transcription directly, which may inhibit *ZCN8*. On the other hand, *ZmCOL3* transcription could have an impact on the circadian clock and down-regulate the clock output *GIGZ1A or GIGZ1B*, thus influence flowering. At the same time, this model also reminds us of the important roles of CCT genes because the majority of genes participating in the maize photoperiod pathway contain a CCT domain.

## Discussion

CCT genes, prevalent in Poaceae, participate in the circadian clock and the photoperiod pathway and have been shown to affect flowering time in *Arabidopsis* and rice. *Ghd7*, the specific heading data regulator component in rice, can delay rice flowering time by repressing *Ehd1* expression and reducing the synthesis of florigen (Xue *et al.*, 2008). *ZmCCT* is the homolog of *Ghd7* and may repress florigen *ZCN8* to delay flowering in maize (Yang *et al.*, 2013). *ZmCCT* is a key gene affecting maize adaptation from tropical to temperate regions. Whether a middle node between *ZmCCT* and *ZCN8* exists in maize flowering pathway remains unclear. *OsCOL4* is a constitutive repressor functioning upstream of *Ehd1* in rice (Lee *et al.*, 2010). The homolog gene in maize, *ZmCOL3*, seems also to act as a repressor in the circadian clock and interacts with *ZmCCT*, causing late flowering by increasing the expression of *ZmCCT*.

Owing to the conservation of the flowering time pathway, it is useful to search for candidate flowering genes and construct flowering time networks by homolog analysis. At the same time, QTL mapping and association analysis also provide abundant flowering time candidate resources. Confidence is increased by combining multiple means of identifying candidate genes. Among the 53 maize CCT genes, 13 were homologous to rice known heading date genes, 28 were located in flowering time QTL regions, and 16 were significantly associated with flowering time. Three genes were found at intersection, including *ZmCOL3*.

*ZmCOL3,* a maize flowering repressor, was thought to function in long day conditions. It may transactivate *ZmCCT* transcription or interfere with circadian clock to inhibit flowering. The loss of one cytosine in its 3’UTR and the presence of a 551bp fragment in the promoter may reduce its transcription and promote maize adaptation from tropical to temperate regions. *ZmCOL3*’s overexpression not only had effects on flowering time, but also influenced many other agronomic traits, including plant height, ear height and internode number. Overexpression of *ZmCOL3* may increase average leaf number, which may increase photosynthesis, thus increasing yield potential, although yield trial tests are still necessary.

Maize was domesticated from teosinte in Mexico in a tropical climate, but currently most maize is planted in temperate regions, including China, which has the largest maize planting area. There are a number of studies exploring the genome wide genetic basis of maize domestication and adaptation (Wright *et al.*, 2005; Hufford *et al.*, 2012). It has been estimated that more than one thousand genes were involved in the domestication and adaptation process (Wright *et al.*, 2005) and few of them seem to overlap (Hufford *et al.*, 2012). Key genes have already been cloned and validated (Doebley *et al.*, 2006). However, few genes have been fully demonstrated to function in the adaptation process. It is hypothesized that transcriptome-level regulatory changes may be a flexible and dynamic way for maize to adapt to environmental changes along its short evolutionary history (Liu *et al.*, 2015). Here, we showed that CCT genes may be the key gene family to affect maize flowering time by regulating the gene expressions and thus help maize rapidly adapt to different environments. It has been shown that many favorable traits, such as disease resistance (Zuo *et al.*, 2015), drought tolerance (Liu *et al.*, 2013; Mao *et al.*, 2015; Wang *et al.*, 2016), and nutritional value (Harjes *et al.*, 2008), are present in the tropical germplasm, and flowering time is one of these. CCT genes and the pathways they are involved in provide clues to our understanding of the underlying molecular mechanism of maize flowering time, thus facilitate maize genetic improvement in the future.

## Supplementary data

**Fig. S1.** Breeding flow of transgenic events

**Fig. S2.** The chromosome position of maize CCT genes

**Fig. S3.** Phylogenetic analysis of CCT genes in four grass species

**Fig. S4.** Candidate genes association mapping results of DT, DA, and ASI in different locations

**Fig. S5.** The Venn diagram of QTLs, candidate genes association mapping, and homology search results

**Fig. S6.** Expression pattern of *ZmCOL3* in 11 tissues combined from 60 different organism parts

**Fig. S7.** The diagram of pCAMBIA3300-UBI: *ZmCOL3* vector and the DNA fragment of *ZmCOL3* used to transform

**Fig. S8.** Transcription of *ZmCOL3* in transgenic lines, controls, receptor HiII, and B73

**Fig. S9.** Expression of *ZmCOL3* in transgenic plants and controls in DHN

**Fig. S10.** Expression of *ZmCOL3* in transgenic plants and controls of RNAi in JL

**Table S1**. Environments used to evaluate association population

**Table S2**. Primers used in this study

**Table S3**. Detailed information of maize CCT genes

**Table S4.** Detailed information of CCT genes in other 3 species

**Table S5**. Maize CCT genes located in flowering time QTL regions

**Table S6.** Candidate genes association mapping results

**Table S7**. Homologs of rice heading date genes in maize

**Table S8.** Phenotype data of transgenic lines

**Table S9.** Alleles of InDel-3296

**Table S10**. Genotype of InDel-551/217 in 317 lines

**Table S11**. The flowering time difference between two InDel types

**Table S12.** Genotype and phenotype of F_2_ population in 3 locations

**Table S13.** DEGs analysis results of circadian clock genes in DHN

## Acknowledgements

This research was supported by the National Key Research and Development Program of China (2016YFD0101003), the National Natural Science Foundation of China (31525017), Huazhong Agricultural University Independent Scientific & Technological Innovation Foundation, Science and Technology Development Plan of Jilin Province (20160520060JH) and Agricultural Science and Technology Innovation Program of Jilin Province (CXGC2017JQ014).

